# Does *Drosophila sechellia* escape parasitoid attack by feeding on a toxic resource?

**DOI:** 10.1101/2020.02.06.937631

**Authors:** Laura Salazar-Jaramillo, Bregje Wertheim

## Abstract

Host shifts can drastically change the selective pressures that animals experience from their environment. *Drosophila sechellia* is a species restricted to the Seychelles islands, where it specialized on the fruit *Morinda citrifolia* (noni). This fruit is known to be toxic to closely related *Drosophila* species, including *D. melanogaster* and *D. simulans*, releasing *D. sechellia* from interspecific competition when breeding on this substrate. Previously, we showed that *D. sechellia* is unable to mount an effective immunological response against wasp attack, while the closely-related species can defend themselves from parasitoid attack by melanotic encapsulation. We hypothesized that this inability constitutes a trait loss due to a reduced risk of parasitoid attack in noni. Here we present a field study aimed to test the hypothesis that specialization on noni has released *D. sechellia* from the antagonistic interaction with its larval parasitoids. Our results from the field survey indicate that *D. sechellia* was found in ripe noni, whereas another *Drosophila* species, *D. malerkotliana*, was present in unripe and rotting stages. Parasitic wasps of the species *Leptopilina boulardi* emerged from rotten noni, where *D. malerkotliana* was the most abundant host. These results indicate that the specialization of *D. sechellia* on noni has indeed drastically altered its ecological interactions, leading to a relaxation in the selection pressure to maintain parasitoid resistance.

## INTRODUCTION

Host shifts are considered to be of major importance in the ecology and evolution of organisms (Nyman, 2010). Adaptation to feeding on novel host-plant species is largely believed to promote speciation and to be a key factor underlying the diversity of insects Ehrlich and Raven (1964); Matsubayashi et al. (2009); Futuyma and Agrawal (2009); Nyman (2010). Among the factors that explain radiaton followed by host shifts are access to new food sources, changes in the competitive dynamics among species and enemy-free spaces (Hardy and Otto, 2014; Janz et al., 2006; Nyman et al., 2007; Feder, 1995). Several studies have focused on the genomic changes associated with these adaptations (Soria-Carrasco et al., 2014; Matzkin, 2012; Feder et al., 2003; Simon et al., 2015). One of the pioneer species used to study the traces left in the genome after a host shift is the specialist species *Drosophila sechellia*. The sequencing of several *Drosophila* species, including the specialist *D. sechellia*, provided a unique opportunity for comparative analysis with respect to the closely related generalist species *D. melanogaster* and *D. simulans* (Clark et al., 2007).

*D. sechellia* is restricted to the Seychelles islands in the Indian Ocean, where it specialized on the fruit *Morinda citrifolia* (commonly known as noni) (Louis and David, 1986; Gerlach, 2009). The noni is toxic to most *Drosophila* species (Farine et al., 1996). A study on the biochemical basis of the toxicity of noni revealed that *D. sechellia* was five to six times more resistant than *D. melanogaster* to one of the toxic compounds, octanoic acid. This toxin is present at high concentration in the ripe stage of the fruit, but less so in the overripe rotten and unripe stages (Legal et al., 1994). In the Seychelles, *D. sechellia* is found abundantly and preferentially on *M. citrifolia* fruits, with a small proportion of adults also found in other substrates (Matute and Ayroles, 2014). Adults of *D. simulans* have also been reported in *M. citrifolia*, but it is not clear whether they are able to breed in this fruit (Matute and Ayroles, 2014). It is believed that resistance of *D. sechellia* to the octanoic acid levels during the highest peak in toxicity, provides this species with a reproductive advantage by being able to access the food source during an earlier time in the fruit’s development (Andrade-López et al., 2017), thus minimizing competition. Genomic changes associated to *D. sechellia*s specialization in the noni have been described for smell and taste receptors (McBride, 2006), as well as genes associated with its resistance to octanoic acid (Jones, 1998; Andrade-López et al., 2017).

In addition to the specialization of traits, host shifts may also lead to trait loss. The change in ecological interactions may alter the selective pressures, resulting either in relaxation of selection for specific traits, or even driving for trait loss when these traits become maladaptive due to altered costs and benefits (Ellers et al., 2012; Brady et al., 2019). This scenario could also apply for *D. sechellia’s* specialization on noni. We hypothesized that its host shift provided it with the protection from the attack by parasitoid wasps, leading to the loss of the immunological resistance against parasitoids (Salazar-Jaramillo et al., 2014).

Parasitoids can constitute a large mortality factor for *Drosophila* species (Janssen et al., 1987; Wertheim et al., 2006). Some species of *Drosophila* can defend themselves after parasitoid attack through an immune response, which is termed melanotic encapsulation. For this, the parasitoid egg in the host larva is detected by the host as a “foreign body”, surrounded with multiple layers of hemocytes (blood cells), and fully melanized. This kills the parasitoid egg, and enables the host to survive the parasitoid attack (Lemaitre and Hoffman, 2007). The differentiation and mobilization of hemocytes is a critical step in this process (Fauverque and Williams, 2011). In *D. melanogaster* three types of differentiated blood cells have been described: 1) plasmatocytes, which perform phagocytosis of bacteria and other small pathogens and are also recruited in the cellular capsules around parasitoid eggs, 2) crystal cells, which store the precursors of the melanin that is deposited on invading pathogens (Pech and Strand, 1996; Williams, 2007) and 3) lamellocytes, which are large, adhesive and flat cells that form the cellular layers around the foreign bodies (e.g., parasitoid eggs) and contain precursors for melanization.

While studying the immune response to parasitoid attack in *Drosophila* species we found (as well as others did before) that *D. sechellia* was unable to defend itself against the infection of the wasp through melanotic encapsulation, while all other tested species of the melanogaster subgroup could, irrespective of the wasp species (Eslin and Prévost, 1998; Schlenke et al., 2007; Salazar-Jaramillo et al., 2014). Other *Drosophila* species outside the melanogaster group also lack the ability to resist parasitoid attack through melanotic encapsulation. In contrast to these species, though, this was not due to the absence of lamellocytes: *D. sechellia* produced lamellocytes in response to parasitoid attack, although in very low concentrations (Eslin and Prévost, 1998; Salazar-Jaramillo et al., 2014).

Two comparative studies, one on genomes and the second on transcriptomes, revealed molecular signatures associated to a loss of resistance. In the comparative genomic study of 12 *Drosophila* genomes, we showed large sequence changes in several of the putative immunity genes uniquely in the genome of *D. sechellia*. In particular, two immune genes showed a potential loss of function sequence variation only in *D. sechellia*: 1) *Tep1*, which facilitates the recognition of pathogens, activation of immune pathways and phagocytosis (Dostálová et al., 2017) and 2) *PPO3*, which is expressed in lamellocytes and contributes to melanization during the encapsulation process (Dudzic et al., 2015). In *Tep1*, we found missing exons; in *PPO3*, we found that the dN/dS ratio was close to unity, suggesting neutral evolution. The disproportionate rate of nucleotide substitution was later shown by another study to correspond to an inactivating mutation due to introduction of a stop codon (Dudzic et al., 2015). When we examined the expression of both genes through qPCR, they were not up-regulated in *D. sechellia* in response to parasitoid attack while they were strongly up-regulated in both *D. melanogaster* and *D. simulans* (Salazar-Jaramillo et al., 2014).

In a comparative transcriptomic study, we further revealed evidence of the failure of a functional immune response in *D. sechellia* to parasitoid attack (Salazar-Jaramillo et al., 2017). Although *D. sechellia* showed upregulation of a few immune genes that react to a general immune challenge (e.g. the homologs of *Mtk, DptB, PGRP-SB1*), it failed to upregulate most of the genes that were upregulated during the cellular immune response against parasitoids in the closely related species *D. melanogaster* and *D. simulans* (Salazar-Jaramillo et al., 2017). These genes included *TepI, PPO3, CG4259, CG4793, TotA* and *Spn88EB*. Figure 1 summarizes the relevant previous genomic and transcriptomic results. Based on these combined results we proposed the hypothesis that *D. sechellia* possesses the “machinery” for melanotic encapsulation, but that it does no longer function properly in response to parasitoid attack (Salazar-Jaramillo et al., 2017). This would signify a case of trait loss.

**Figure 1.**
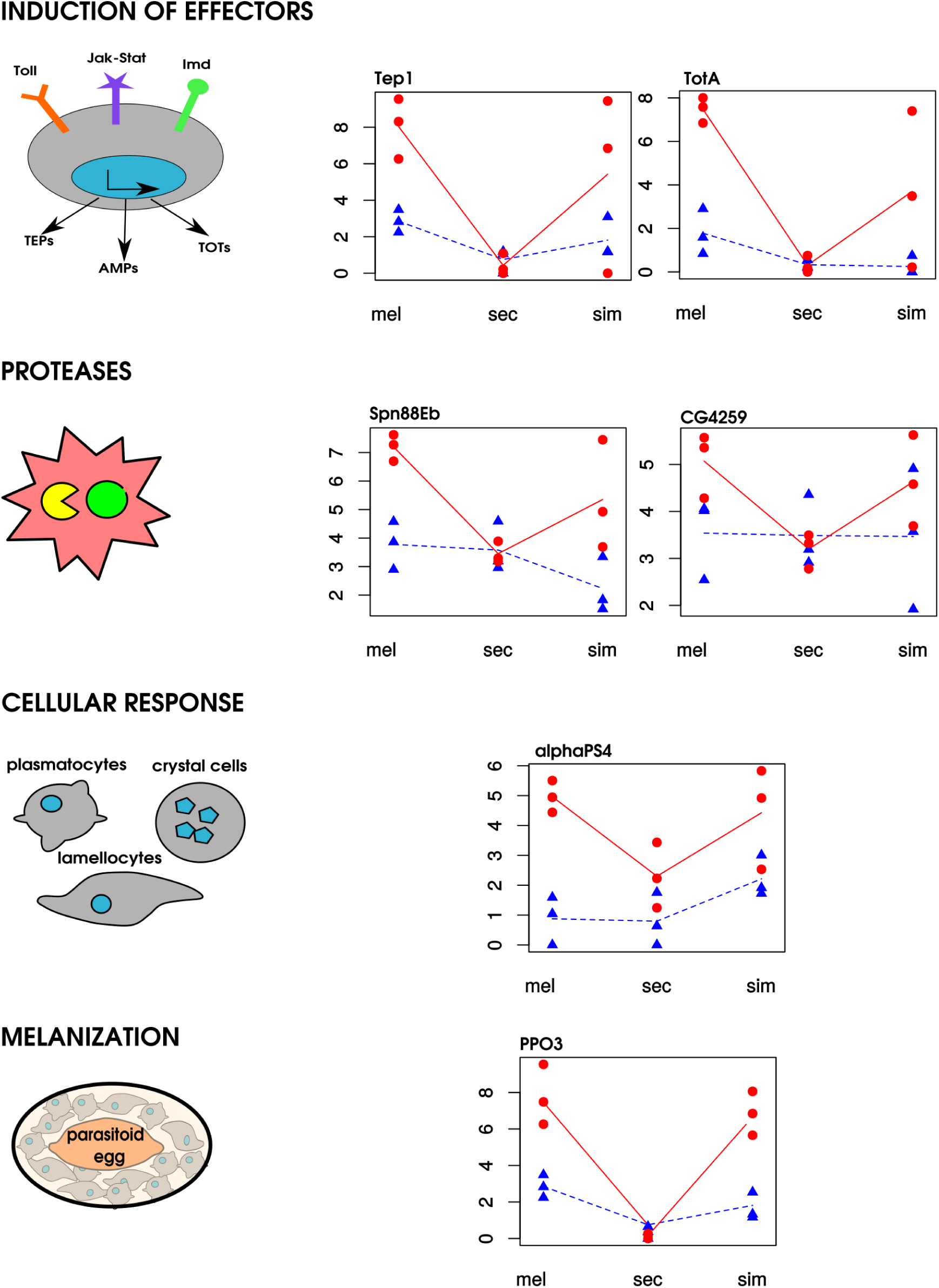
Differential expression of genes that show up-regulation in *D. melanogaster* and *D. simulans* after parasitoid attack but not in *D. sechellia* (right column), with their respective functional annotation during the immune response (left column). Red cicrcles show expression data (Log2 Counts per million) upon wasp parasitization and blue triangles the expression of the unparasitized controls. Each biological replicate (n=3) is the expression of 5 pooled larvae at 5h or 50h after parasitoid attack. Data come from the RNA study in Salazar-Jaramillo et al. (2017), except for PPO3, which is based on the RT-qPCR assay in Salazar-Jaramillo et al. (2014) (due to gene model error in FlyBase that fuses incomplete versions of PPO3 and a neighboring gene in *D. sechellia* as pointed out in Dudzic et al. (2015)). The connecting lines is shown to emphasize the trend

The question we address in this manuscript is whether the observed trait loss for parasitoid resistance in *D. sechellia* may be the consequence of its host shift to noni. Our hypothesis is that this new host provides an enemy-free space. To test this hypothesis, we need to establish the occurrence or absence of parasitoid wasps in noni fruit. Here we present the results of a field study in Couisin, one of the Seychelles Islands to test whether *D. sechellia* flies that breed and feed on noni would experience parasitism by parasitoid wasps. We collected noni fruits at different levels of maturity, and reared out insects that developed on these fruits. Cousin has the status of Special Reserve for nature conservation and due to its conservation status, it is not allowed to extract live material, which limited our sample sizes considerably. Therefore the field study may not be conclusive on its own, but it is compelling in the light of the multiple lines of evidence.

## MATERIALS AND METHODS

### Field study

We did a survey of Drosophilid flies and parasitoid wasps occurring on wild noni fruit on Cousin island in the Seychelles. Cousin Island is a nature reserve characterized by the presence of indigenous and endemic forest (mixed Pisonia, Noni and Ochrosia). We collected noni fruit that had fallen of the plant, at different stages of maturity. A total of 28 noni fruit were collected, ranging from unripe, to ripe, to rotten. Collected unripe fruits were allowed to mature and considered “ripe” after one week and “rotten” after two weeks of collection. The fruits were placed in plastic containers to capture larvae that would leave the fruit for pupation. The containers and fruits were left open at the site of collection to enable further oviposition by insects, for a period varying from 1-5 days depending on the stage of maturity of the noni fruit. Thereafter, the containers were closed with a piece of gauze to ensure that all insects that emerged from the fruit could be retrieved. The containers were brought to the field station and checked regularly for any emerging insects. Emerged adult insects were collected and preserved in 70% alcohol for taxonomic identification.

In order to assess whether other *Drosophila* species were present on Cousin Island we also placed out 21 alternative fruit baits containing a substrate of North Carolina instant medium and a layer of either banana, papaya or noni. Each set of three fruit types were placed in 7 different locations across the island and left during 24 hours to allow flies to lay eggs. Emerged larvae were transferred to fresh medium and emerged adults stored in 70% alcohol for taxonomic identification.

### Species identification

*Drosophilidae* species were identified by Prof Marie-Louise Cariou using morphological traits (genitalia of males allowed the identification at the species level) (Lachaise et al., 2008). Emerged wasps were identified through morphological characters (antennae, wing venation, scutellum and shape of the scutellar cup) and through sequencing of a Cytochrome Oxidase I fragment (Lue et al., 2016)

## RESULTS

We found two species of *Drosophila, D. sechellia* and *D. malerkotliana*, in both types of samples, the collected noni and the fruit baits. Our characterization of the Drosophilid fruit fly and parasitoid wasp community that developed from the noni revealed that the stage of maturity was decisive for the number and type of species that emerged from it (Table 1).

**Table 1.**
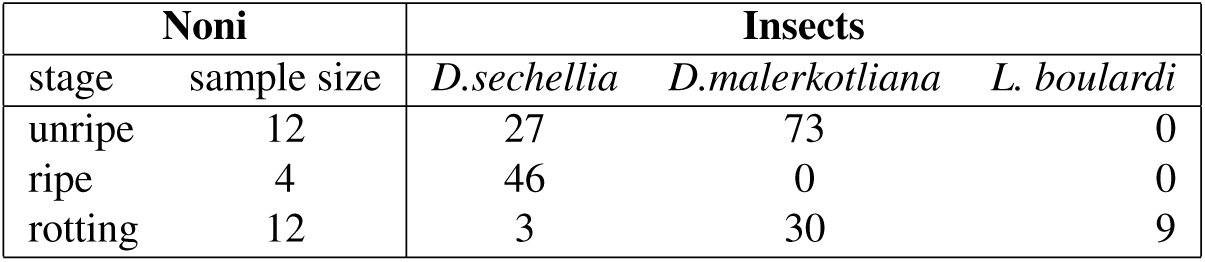
Summary of results. Sample sizes of noni with their stages and emerged insects.

Only *D. sechellia* was found in ripe noni, whereas another *Drosophila* species, *D. malerkotliana*, was abundant in unripe noni and also present in the rotting stage but completely absent from the ripe stage. Parasitic wasps emerged from two rotten noni fruit, around seven weeks after the collection of the fruit. The morphological and molecular analysis identified the wasp species as *Leptopilina boulardi*. The typical developmental time of these parasitoids at 20 − 25°C is approximately 4 weeks, meaning that they had infested their host during the rotting stage of noni, when *D. malerkotliana* was the predominant species. No wasps emerged from noni fruits that were collected during the ripe phase when *D. sechellia* was dominant.

## DISCUSSION

Inspired by evidence from phenotypic, genomic and transcriptomic studies, we hypothesized that *D. sechellia*’s specialization on noni fruit may have protected it from the infection by parasitoid wasps. This could cause the relaxation in the selection pressure to maintain parasitoid resistance, and thereby lead to trait loss. The evidence included the lack of resistance against parasitic wasps despite producing lamellocytes, and the changes in the sequence and expression found in genes involved in the immune response against parasitoid attack compared to closely related species.

Fundamental to this hypothesis is the occurrence or absence of parasitoid wasps developing in *D. sechellia* that feeds and breeds on noni. In our field study on the Seychelles, we did in fact find a small number of *L. boulardi* parasitoids emerging from noni fruit, but only in the rotting stages when *D. sechellia* was very rare and *D. malerkotliana* was abundant. In the (small number of) ripe noni fruits we collected, from which only *D. sechellia* fruit flies were reared, no parasitoids were recorded. This supports our hypothesis that the specialization of *D. sechellia* on ripe noni fruits, containing high concentrations of octanoic acid, may provide a parasitoid-free space, which may have led to the loss of the immunological defence against parasitoids.

It is important to mention that the policy of collection in the Seychelles is very restrictive, particularly concerning live material, which cannot be extracted from the Islands. This imposed strong limitations to the sample sizes, because all the data had to be obtained *in situ*. In addition, many samples were lost due to uncontrolled conditions (e.g., rain washing out some of our collected fruits, and animal invasions raiding our noni samples while they were still out in the field). Despite these shortcomings, our survey confirmed that *D. sechellia* was most abundant during the ripe stage of noni (e.g., in fruits approx. 1 week after falling to the ground), as predicted by its tolerance to the toxin in the early stages (Legal et al., 1994). Figure 2 summarizes the hypothetical scenario of insects developing in noni. At earlier maturation stages (less than a week) and later (more than 2 weeks) stages of decomposition, we found one more species of *Drosophila, D. malerkotliana*.

**Figure 2.**
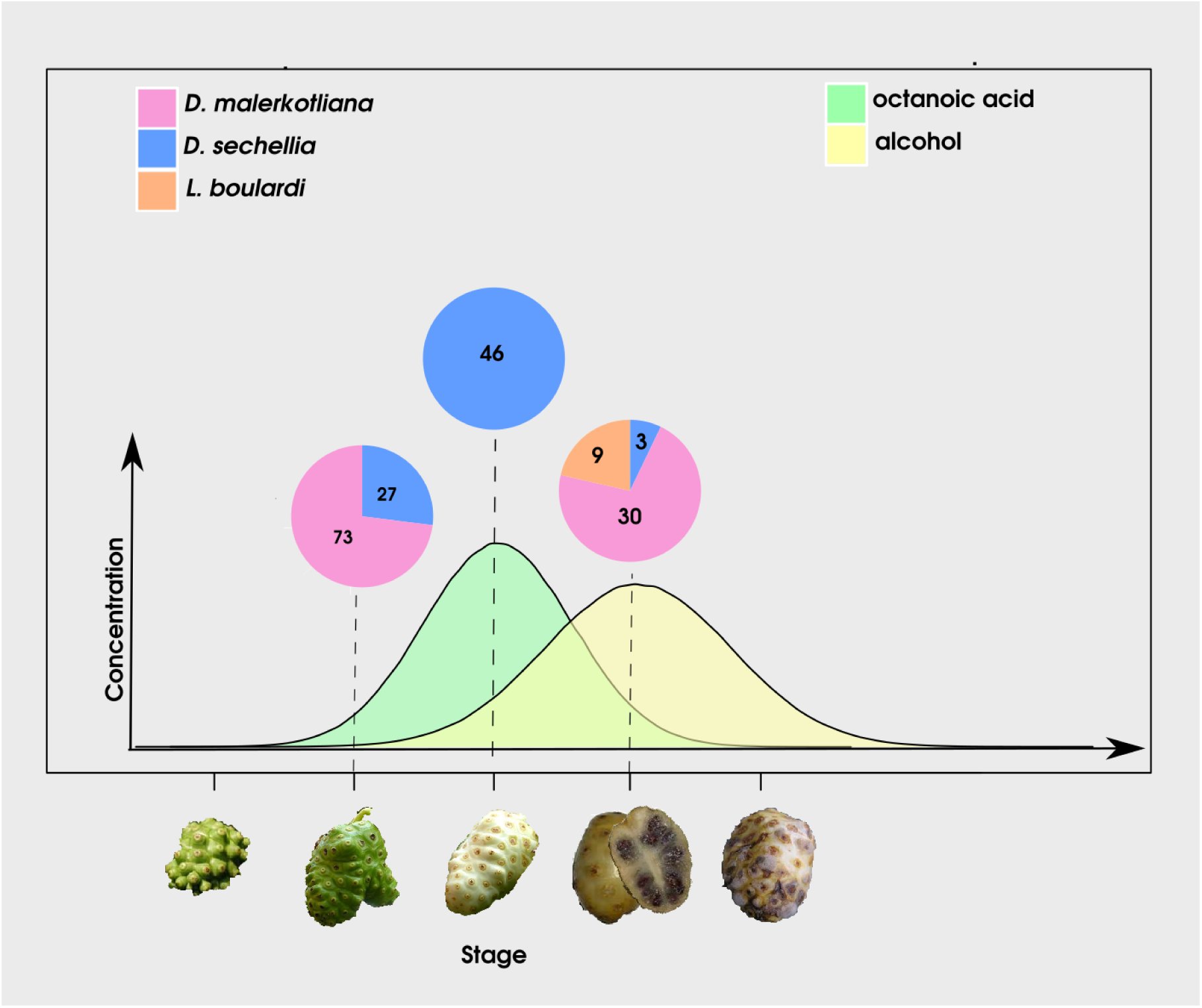
Number of insects emerging from *Morinda citrifolia* at different stages of maturity. A hypothetical concentration curve of alcohol and octanoic acid in the fruit is included based on (Andrade López, 2015)

Parasitic wasps from the species *L. boulardi* were only recovered in a late stage of decomposition, which mostly contained larvae from *D. malerkotliana*. It is, however, well known that parasitic wasps, and particularly *L. boulardi*, can develop in *D. sechellia* (Schlenke et al., 2007; Lee et al., 2009). Although not conclusive, this result provides support for the hypothesis that the toxins in the ripe stage of noni protect *D. sechellia* from parasitic wasps, and that this protection released the selection pressure to maintain the mechanism of melanotic encapsulation.

Parasitoids are considered an important selection force due to the heavy mortality they can inflict on other insects, thus release from this enemy could have played an important role in accelerating the diversification of *D. sechellia*. It remains to be tested what the effect of the ripe noni is on the parasitic wasps to understand whether their absence was due to lethality or avoidance. An artificial incubation experiment with another parasitoid species (*Asobara citr*i) does suggest that parasitoids can indeed experience lethality and refrain from parasitizing in the presence of noni: 24 h exposure of the female parasitoid in petri-dishes to noni extract (“Hawaiian Health Ohana”) with 25 *D. melanogaster* or *D. simulans* larvae resulted in 50% mortality of the female parasitoids and low rates of parasitism (*<* 20% on average, n=12), while a similar exposure to petri-dishes with 25 *D. melanogaster* or *D. simulans* larvae on yeast resulted in 0% mortality and on average *>* 90% parasitism (Laura Salazar Jaramillo, unpublished data available in the github repository: https://github.com/lauraalazar/Dsechellia-parasitoid)

*D. malerkotliana* is an invasive species, which has been found to be a strong competitor of *D. sechellia* (Lachaise et al., 2008). A study showed that this species (together with other species from the ananassae group) are able to encapsulate and kill parasitoid eggs without melanization, by means of gigantic cells with filamentous projections and multiple nuclei (named multinucleated giant hemocyte, MGH). These cells share some properties with lamellocytes, such as the ability to encapsulate foreign objects, but differ considerably in their morphology and function (Márkus et al., 2015). It is thus intriguing whether *D. malerkotliana*’s ability to resist parasitoid wasps helped it to invade the decaying noni fruits, where *Leptopilina* parasitoids have been found, thus filling a niche in exploiting rotting stages of the noni.

## CONCLUSION

Collectively, our data indicate that a consequence of *D. sechellia*’ s ecological shift to noni may have protected it from parasitic wasps. This, in turn, generated a relaxation of the selective pressure to maintain the functionality of genes that are essential in the immune respones against the parasitoid wasp infection, leading to the loss of this trait.

## ACKNOWLEDGMENTS

We would like to thank Marie-Louise Cariou for the taxonomic identification of the *Drosophila species* and Fabrice Vavre for conduting the amplification of the COI fragment of the parasitic wasps. To the NGO *Nature Seychelles* and the team of volunteers on Cousin island for supporting the field work, in particular to Alex Underwood for assisting with the collection sites. The hypothetical curves of octanoic acid and alcohol in Figure 2 are based on Andrade-Lopez MSc thesis.

## REFERENCES

Andrade-López, J., Lanno, S. M., J.M. Auerbach, E.C. Moskowitz, L. S., Wittkopp, P., and Coolon, J. (2017). Genetic basis of octanoic acid resistance in *Drosophila sechellia*: functional analysis of a fine-mapped region. Molecular Ecology, 26:1148–1160.

Andrade López, J. M. (2015). Genetic basis of octanoic acid resistance in Drosophila sechellia: functional analysis of a fine-mapped region.

Brady, S. P., Bolnick, D. I., Angert, A. L., Gonzalez, A., Barrett, R. D., Crispo, E., Derry, A. M., Eckert, C. G., Fraser, D. J., Fussmann, G. F., Guichard, F., Lamy, T., McAdam, A. G., Newman, A. E., Paccard, A., Rolshausen, G., Simons, A. M., and Hendry, A. P. (2019). Causes of maladaptation. Evolutionary Applications, 12(7):1229–1242.

Clark, A. G., Eisen, M. B., Smith, D. R., and *Drosophila* 12 Genomes Consortium (2007). Evolution of genes and genomes on the Drosophila phylogeny. Nature, 450(7167):203–218.

Dostálová, A., Rommelaere, S., Poidevin, M., and Lemaitre, B. (2017). Thioester-containing proteins regulate the toll pathway and play a role in *Drosophila* defence against microbial pathogens and parasitoid wasps. BMC Biology, 15:79.

Dudzic, J. P., Kondo, S., Ueda, R., Bergman, C. M., and Lemaitre, B. (2015). *Drosophila* innate immunity: regional and functional specialization of prophenoloxidases. BMC Biology, 13(1):1–16.

Ehrlich, P. R. and Raven, P. H. (1964). Butterflies and plants: a study in coevolution. Evolution, 18:586–608.

Ellers, J., Toby Kiers, E., Currie, C. R., McDonald, B. R., and Visser, B. (2012). Ecological interactions drive evolutionary loss of traits. Ecology Letters, 15(10):1071–1082.

Eslin, P. and Prévost, G. (1998). Hemocyte load and immune resistance to *Asobara tabida* are correlated in species of the *Drosophila melanogaster* subgroup. Journal of Insect Physiology, 44:807–816.

Farine, J., Legal, L., Moreteau, B., and Le Quere, J. (1996). Volatile components of ripe fruits of *Morinda citrifolia* and their effects on *Drosophila*. Phytochemestry, 41(2):433–438.

Fauverque, M.-O. and Williams, M. J. (2011). *Drosophila* cellular immunity: a story of migration and adhesion. Journal of Cell Science, 124:1373–1382.

Feder, J. L. (1995). The effects of parasitoids on sympatric host races of *Rhagoletis pomonella* (Diptera: Tephritidae). Ecology, 76:801–813.

Feder, J. L., Berlocher, S. H., Roethele, J. B., Dambroski, H., Smith, J. J., Perry, W. L., Gavrilovic, V., Filchak, K. E., Rull, J., and Aluja, M. (2003). Allopatric genetic origins for sympatric host-plant shifts and race formation in *Rhagoletis*. Proceedings of the National Academy of Sciences, 100(18):10314–10319.

Futuyma, D. J. and Agrawal, A. (2009). Evolutionary history and species interactions. Proceedings of the National Academy of Sciences, 106:18043–18044.

Gerlach, J., editor (2009). The Diptera of the Seychelles. Pensoft, Sofia, Bulgaria.

Hardy, N. B. and Otto, S. P. (2014). Specialization and generalization in the diversification of phytophagous insects: tests of the musical chairs and oscillation hypotheses. Proceedings of the Royal Society B, 281:1795.

Janssen, A., Driessen, M., De Haan, M., and Roodbol, N. (1987). The impact of parasitoids on natural populations of temperate woodland *Drosophila*. Netherlands Journal of Zoology, 38(1):61–73.

Janz, N., Nylin, S., and Wahlberg, N. (2006). Diversity begets diversity: host expansions and the diversification of plant-feeding insects. BMC Evolutionary Biology, 6.

Jones, C. (1998). The genetic basis of *Drosophila sechellia* resistance to a host plant toxin. Genetics, 149:1899–1908.

Lachaise, D., Gerlach, J., Matyot, P., Legrand, D., Montchamp, C., and Cariou, M. (2008). Drosophilidae of Seychelles: biogeography, ecology and conservation status. Phelsuma, 15:19.

Lee, M. J., Kalamarz, M. E., Paddibhatla, I., Small, C., Rajwani, R., and Govind, S. (2009). Virulence factors and strategies of *Leptopilina* spp.: Selective responses in *Drosophila* hosts. Advances in Parasitology, 70:123–145.

Legal, L., Chappe, B., and Jallon, J. (1994). Molecular basis of *Morinda citriflora* toxicity on *Drosophila*. Journal of Chemical Ecology, 20:1931–1943.

Lemaitre, B. and Hoffman, J. (2007). The host defence of *Drosophila melanogaster*. Annual Review in Immunology, 25:697–743.

Louis, J. and David, J. (1986). Ecological specialization in the *Drosophila melanogaster* species subgroup: a case-study of *Drosophila sechellia*. Acta Oecologica, 7:215–230.

Lue, C. H., Driskel, A., Leips, J., and Mathew, B. (2016). Review of the genus *Leptopilina* (Hymenoptera, Cynipoidea, Figitidae, Eucoilinae) from the Eastern United States, including three newly described species. Journal of Hymenoptera Research, 53:1229–1242.

Márkus, R., Lerner, Z., Honti, V., Csordás, G., J, Z., Cinege, G., Párducz, A., Lukacsovich, T., Kurucz, E., and Andó, I. (2015). Multinucleated giant hemocytes are effector cells in cell-mediated immune responses of *Drosophila*. Journal of Innate Immunity, 7(4):340–353.

Matsubayashi, K. W., Ohshima, I., and Nosil, P. (2009). Ecological speciation in phytophagous insects. Entomologia Experimentalis et Applicata, 134:1–27.

Matute, D. and Ayroles, J. (2014). Hybridization occurs between *Drosophila simulans* and *D. sechellia* in the seychelles archipelago. Journal of Evolutionary Biology, 27:1057–68.

Matzkin, L. (2012). Population transcriptomics of cactus host shifts in *Drosophila mojavensis*. Molecular Ecology, 21:2428–2439.

McBride, C. S. (2006). Rapid evolution of smell and taste receptor genes during host specialization in *Drosophila sechellia*. Proceedings of the National Academy of Sciences, 104:4996–5001.

Nyman, T. (2010). To speciate, or not to speciate? Resource heterogeneity, the subjectivity of similarity, and the macroevolutionary consequences of niche-width shifts in plant-feeding insects. Biological Reviews, 85:393–411.

Nyman, T., Bokma, F., and Kopelke, J.-P. (2007). Reciprocal diversification in a complex plant-herbivore-parasitoid food web. BMC Biology, 6(49).

Pech, L. L. and Strand, M. R. (1996). Granular cells are required for encapsulation of foreign targets by insect haemocytes. Journal of Cell Science, 109:2053–2060.

Salazar-Jaramillo, L., Jalvingh, K. M., de Haan, A., Buermans, K. K. H., and Wertheim, B. (2017). Inter- and intra-species variation in genome-wide gene expression of *Drosophila* in response to parasitoid wasp attack. BMC Genomics, 18:331.

Salazar-Jaramillo, L., Paspati, A., van de Zande, L., Vermeulen, C. J., Schwander, T., and Wertheim, B. (2014). Evolution of a cellular immune response in *Drosophila*: a phenotypic and genomic comparative analysis. Genome Biology and Evolution, 6:273–289.

Schlenke, T. A., Morales, J., Govind, S., and Clark, A. G. (2007). Constrasting infection strategies in generalist and specialist wasp parasitoids of *Drosophila melanogaster*. PLoS Pathogens, 3:e158.

Simon, J.-C., d’Alençon, E., Guy, E., Jacquin-Joly, E., Jaquiéry, J., Nouhaud, P., Peccoud, J., Sugio, A., and Streiff, R. (2015). Genomics of adaptation to host-plants in herbivorous insects. Briefings in Functional Genomics, 14(6):413–423.

Soria-Carrasco, V., Gompert, Z., Comeault, A. A., Farkas, T. E., Parchman, T. L., Johnston, J. S., Buerkle, C. A., Feder, J. L., Bast, J., Schwander, T., Egan, S. P., Crespi, B. J., and Nosil, P. (2014). Stick insect genomes reveal natural selection’s role in parallel speciation. Science, 344(738).

Wertheim, B., Allemand, R., Vet, L., and Dicke, M. (2006). Effect of aggregation pheromone on individual behaviour and food web interactions: field study on *Drosophila*. Ecological Entomology, 31(3):216–226.

Williams, M. J. (2007). *Drosophila* hemopoiesis and cellular immunity. The Journal of Immunology, 178:4711–4716.

